# Propagation of human respiratory syncytial virus in cells derived from the black flying fox (*Pteropus alecto*)

**DOI:** 10.1101/2023.09.03.556015

**Authors:** Timothy Tan, Boon Huan Tan, Richard Sugrue

## Abstract

The propagation of human respiratory syncytial virus (hRSV) was evaluated in the *Pteropus alecto* kidney (PaKi) cell line. At 20 hrs post-infection, immunoblotting of hRSV-infected PaKi cell lysates with anti-G, anti-N, anti-P and anti-M2-1 indicated expression of the respective virus proteins of the correct size. The hRSV-infected PaKi cell were also stained using anti-F, anti-G, anti-N, anti-P and anti-M2-1 and imaged using immunofluorescence microscopy, which confirmed high levels of virus infection, and the presence of numerous virus filaments and virus-induced inclusion bodies. PaKi cell monolayers also supported multiple cycle infection when hRSV was used to infect PaKi cells using a low multiplicity of infection. These data indicate that prior adaptation of hRSV was not required for its propagation in the PaKi cell line, and suggests that PaKi cell line is a suitable cell model system with which to examine virus-host interactions involving RSV infection in fruit bats.

## Introduction

Human respiratory syncytial virus (hRSV) is currently the most important viral cause of lower respiratory tract infection in young children and neonates, leading to high levels of mortality and morbidity [1]. Although RSV was first identified in the 1950s [2, 3], hRSV was detected in lung biopsies taken from individuals in the 1930s and 1940s [4], suggesting that RSV was responsible for human infection prior to its first formal description. Therefore, the precise time of the appearance of hRSV in the human population is uncertain. In this context, strains of RSV infect a range of different mammals (e.g. bovine RSV and cattle) and sheep (ovine RSV) giving rise to similar symptoms to that exhibited by humans that are infected with hRSV strains [5]. The presence of closely related RSV strains in different mammalian hosts also indicates that RSV can adapt to different animal species, but it is unclear to what extent these closely-related animal viruses play in the emergence of new hRSV stains. In this context, the zoonotic origins of other paramyxoviruses that are commonly associated with human disease have been proposed [6, 7], suggesting the possibility that similarly to other paramyxovirus, hRSV may also have had a zoonotic origin.

Although bats play an important and essential role in human habitat sustainability (e.g., in pollination and insect control), they are also established reservoirs for several clinically important emerging paramyxoviruses that cause serious infection in humans [8]. Frequent contact between bats and humans does not normally occur in the natural environment, but on rare occasions direct or indirect contact between humans and bats can lead to the zoonotic transmission of these viruses to humans. In this context, Drexler and colleagues performed a global surveillance using partial virus genomic analysis and detected the presence of several different paramyxoviruses in a variety of different bat species. This study identified several paramyxoviruses in fruit bats (*pteropodidae* species) that were closely related to established human (hRSV) and bovine (bRSV) RSV strains [9]. The relationship between these RSV-like virus strains in bats and hRSV was supported by the cross-reactivity of fruit bat serum with RSV-infected cells, but a detailed evaluation of the evolutionary relatedness of the RSV-like viruses identified in bats and the circulating human and bovine RSV strains was not presented. Therefore, the significance of these bat-associated viruses in the context of RSV epidemiology and the emergence of novel RSV strains that spillover into humans and cattle remain uncertain. Many viruses that spread to humans via a zoonotic route can replicate in cells derived from tissue of both the animal reservoir and human host. A first step to determine if RSV replication is permissive in fruit bats would be to determine if the virus can be propagated in cells derived from fruit bat tissue without prior cell-adaptation of the virus. The *Pteropus alecto* kidney (PaKi) cell line was derived from the fruit bat *Pteropus alecto*, and this cell line supports the replication of several emerging paramyxoviruses whose replication in *Pteropus alecto* has been established [10-12]. The PaKi cell line therefore represents a convenient cell model system with which to initially examine the potential for RSV infection in *Pteropus* bats (e.g., *P. alecto*), and we report the first study to evaluate the propagation of hRSV in the PaKi cell line.

## Results and Discussion

In permissive cells infected with hRSV two distinct virus structures are formed, which are referred to as the virus filaments and inclusion bodies. The virus filaments are the mature form of the virus particles that assembles on the cell surface as filamentous projections [13]. They are surrounded by a lipid envelope in which the attachment (G) protein and fusion (F) protein are inserted, and the virus filaments mediate virus transmission between cells [14, 15]. The virus genomic RNA (vRNA) interacts with the virus encoded nucleocapsid (N) protein, phosphoprotein (P protein), the M2-1 protein and the large (L) protein to form the the virus nucleocapsid (NC), and virus polymerase activity is associated the nucleocapsid. In hRSV-infected cells the nucleocapsid-associated proteins are packaged into the virus filaments and they also accumulate in the cytoplasm as cytoplasmic inclusion bodies.[13-16].

PaKi cells were either mock-infected or infected with hRSV using a multiplicity of infection (moi) of 5, and at 20 hrs post-infection (hpi) cell lysates were prepared. The expression of the G protein, N protein, P protein and M2-1 proteins was examined in hRSV-infected PaKi cells by immunoblotting of the cell lysates using anti-G, anti-N, anti-P and anti-M2-1 respectively (Fig. 1). Protein species of the expected size were detected with each antibody, which confirmed the expression of these hRSV proteins in the RSV-infected PaKi cells. The 100 kDa G protein was the major form of the G protein detected in PaKi cells, and it was similar in size to the G protein detected in human cell lines infected with hRSV. This suggested that similar post-translational processing of the G protein by O-linked and N-linked glycosylation also occurred in PaKi cells. In a parallel analysis, PaKi cells were either mock-infected or infected with hRSV using a multiplicity of infection (moi) of 5 and at 20 hrs post-infection (hpi) the cells were stained using anti-G, anti-F (recognizes the F protein) and anti-anti-N, anti-P and anti-M2-1 and examined by immunofluorescence (IF) microscopy (Fig. 2A). This indicated that in mock-infected cells no staining was observed, while in virus-infected cells greater than 95% of the cells showed staining with each of the virus-specific antibodies.

**Fig. 1.**
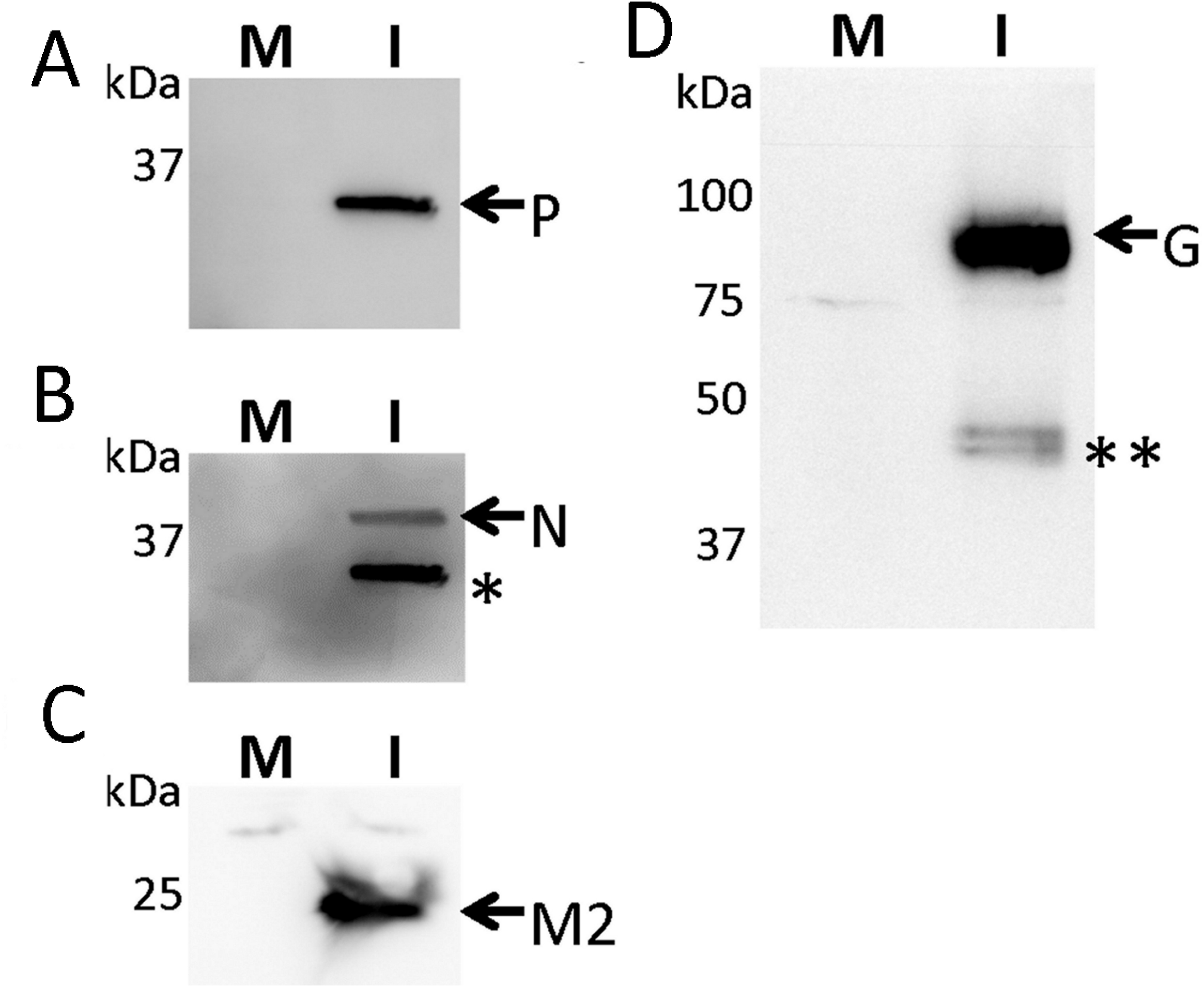
Expression of the RSV structural proteins in RSV-infected PaKi cells. The PaKi cells were mock-infected and RSV-infected using a multiplicity of infection of 5 and at 20 hrs post-infection cell lysates were prepared from the mock-infected (M) and RSV-infected (I) cells and immunoblotted using **(A)** anti-P, **(B)** and anti-N, **(C)** anti-M2-1 and **(D)** anti-G. In **(B)** the membrane originally probed using anti-P (as in **(A))** was then re-probed using anti-N, and the corresponding anti-P labeled band is highlighted (*). **(D)** In immunoblotting using anti-G, the mature glycosylated G protein (black arrow) and the smaller (**) N-linked glycosylated anti-G protein species are indicated.

**Fig. 2.**
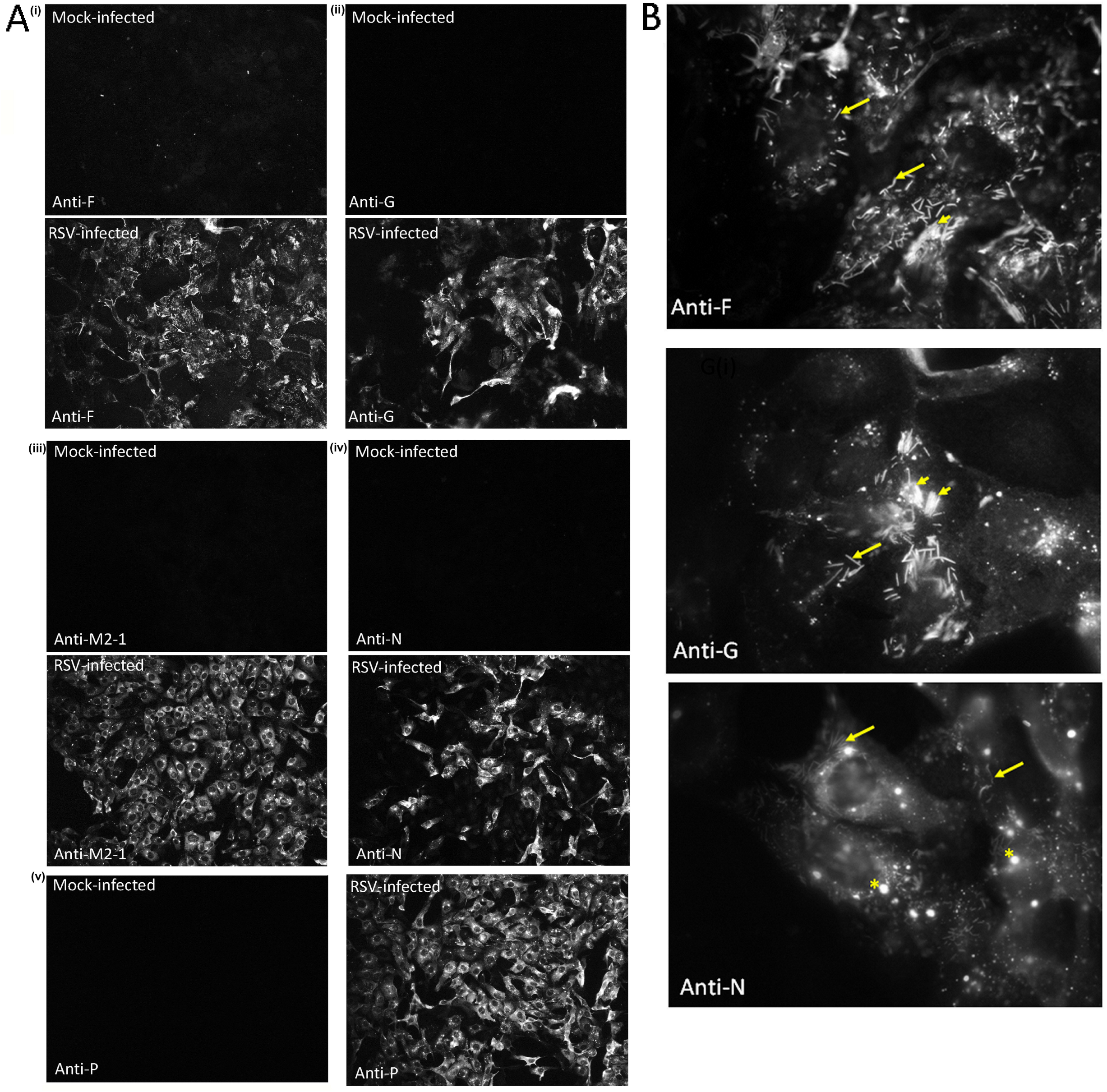
The PaKi cell line is susceptible to infection with RSV. The PaKi cells were mock-infected and RSV-infected using a multiplicity of infection of 5. At 20 hrs post-infection the cells were **(A)** stained using (i) anti-F, (ii) anti-G, (iii) anti-M2-1, (iv) anti-N and (v) anti-P, and the stained cells examined by immunofluorescence microscopy (objective x20 magnification). **(B)** The RSV-infected PaKi cells singly stained using anti-F, anti-G and anti-N were imaged using IF microscopy (objective x100 magnification (oil)). The presence of individual virus filaments (long arrow), inclusion bodies (*) and clusters of virus filaments (short arrows) are highlighted.

In permissive cells that are infected with hRSV the formation of the virus filaments and inclusion bodies are characteristic feature of a productive virus infection. Their presence in hRSV-infected PaKi cells would be expected if these cells were able to support the propagation of hRSV. The hRSV-infected PaKi cells were therefore examined to determine if the virus filaments and inclusion bodies form in hRSV-infected PaKi cells. Cells stained with anti-F, anti-G and anti-N and examined in greater detail by using IF microscopy (Fig. 2B). Staining with each of the three antibodies revealed the presence of a filamentous staining pattern on the surface of the RSV-infected PaKi cells that was consistent with the presence of virus filaments. A high density of virus filaments on the surface of individually infected cells was noted, and in some cases the virus filaments appeared to form in tight clusters on the cell surface. Staining with the anti-N antibody also revealed the presence of anti-N-stained cytoplasmic inclusion bodies in the RSV-infected PaKi cells. Collectively these data indicated that mature hRSV particles were able to form on the surface of hRSV-infected PaKi cells, indicating that PaKi cells were permissive for hRSV infection.

These data indicated that PaKi cells were able to support hRSV replication, and this further suggested that hRSV could be propagated in PaKi cell line and this was examined further. We have previously described a multiple cycle infection model where cell monolayers are infected at low moi (typically between 0.01 and 0.0001), and the spread of hRSV infection in the monolayers examined over extended time periods [17]. The previous study indicated that hRSV infection remained largely cell-associated, and that the spread of infection was due to localized transmission of infectious progeny virus in the monolayer. PaKi cell monolayers that were approximately 95% confluent were infected with hRSV using a moi corresponding to 5, 0.05, 0.005 and 0.001, and at 24 hpi the cells were stained using anti-RSV and examined by IF microscopy (Fig. 3). At a moi of 5, greater than 95% of the cells were infected, while progressively less cells were infected as the moi was decreased. Using a moi of 0.001 approximately 1% of the cells in the monolayer was infected, and this moi was employed in subsequent experiments involving multiple-cycle infection in PaKi cell monolayers. PaKi cell monolayers were mock-infected or infected with hRSV and at between 1 and 4 days-post infection (dpi) the cells were stained using either anti-P (Fig. 4A) or anti-F (Fig. 4B) and examined by IF microscopy. At 1 dpi we noted the predominance of single virus-infected cells with an average cell cluster of 1.0±0.3 cells per cluster, while at 3 dpi the presence of clearly defined infected cell clusters consisting of 24.3±8.7 cells per cluster. At 4 dpi the appearance of infected cell clusters containing greater than 50 cells per cluster was noted in the PaKi cell monolayers stained with either antibody. These data indicated that while individual cells are infected early in infection, as time progresses these single infected cells gave rise to individual clusters of infected cell. This was consistent with localised spread of infection in the PaKi cell monolayer, consistent with our previous observations in different permissive cell lines [17]. The appearance of the infected cell cluster was consistent with a productive infection in the PaKi cell monolayers.

**Fig. 3.**
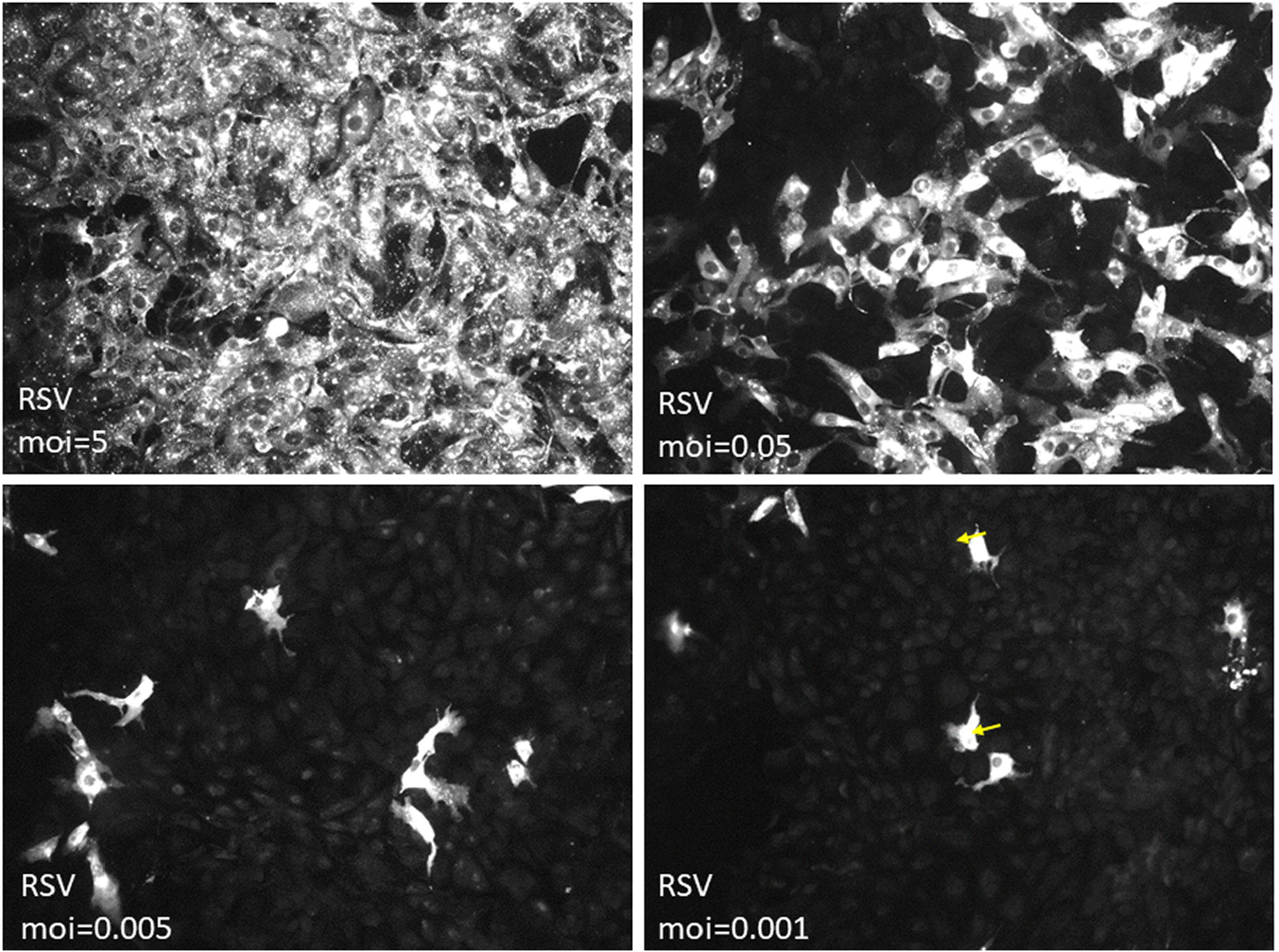
RSV infection PaKi cell monolayers using a low multiplicity of infection. PaKi cell monolayers were infected with RSV using a multiplicity of infection of 5, 0.05, 0.005 and 0.001. At 20 hrs post-infection the cells were stained using anti-P, and the cells examined by immunofluorescence (IF) microscopy (objective x20 magnification).

**Fig. 4.**
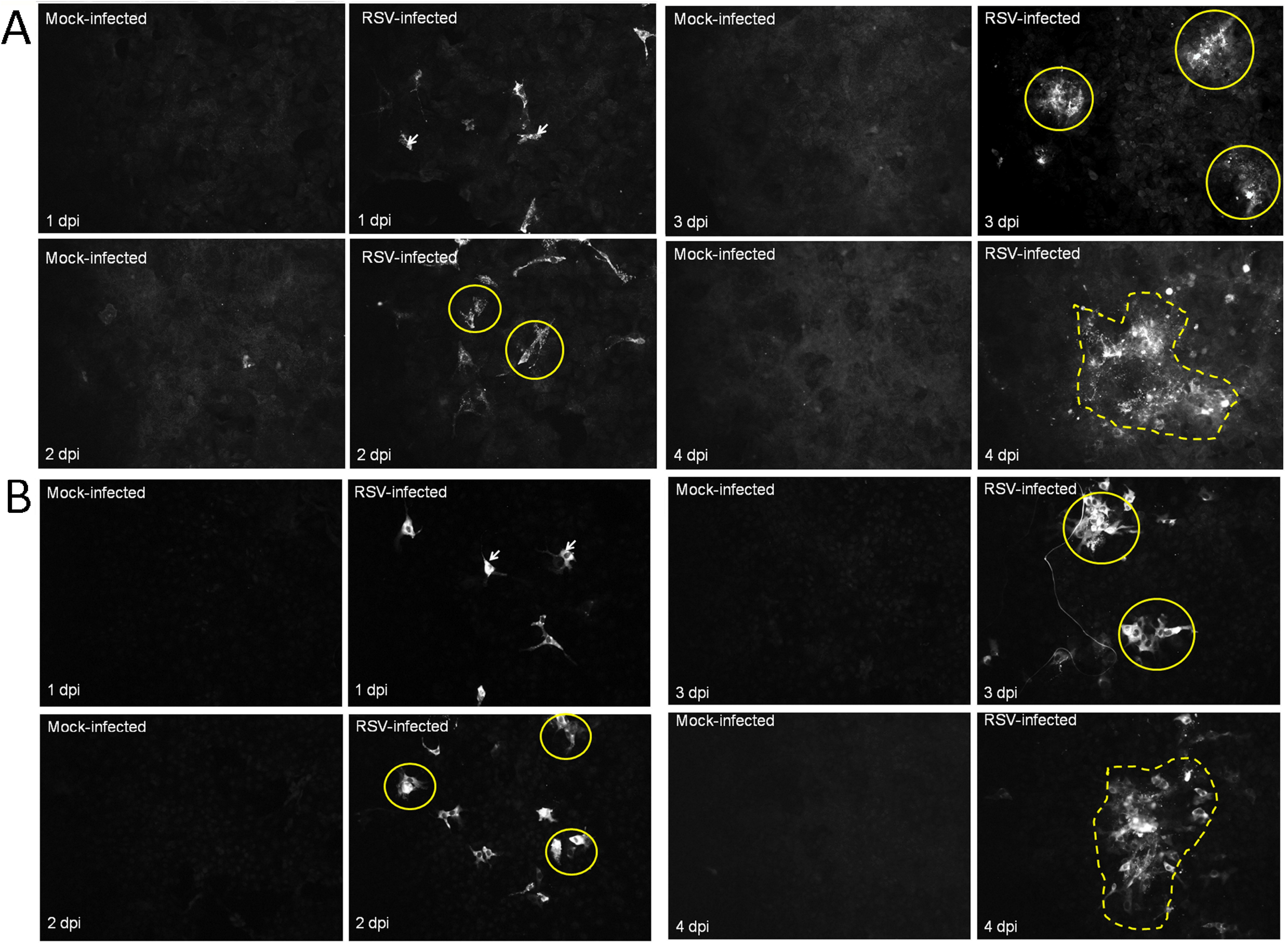
Evidence for multiple cycle RSV transmission in PaKi cell monolayers. PaKi cell monolayers were either mock-infected or infected with RSV using a multiplicity of infection of 0.001 and at 1, 2, 3, and 4 days-post infection the cell monolayers were stained using **(A)** anti-G and **(B)** anti-P. The stained cell monolayers were examined by immunofluorescence microscopy (objective x20 magnification). Individual infected cells at 1 dpi (yellow arrows), infected cell clusters at 2 and 3 dpi (yellow circles) and the larger infected cell clusters (broken yellow lines) at 4 dpi are highlighted.

PaKi cell monolayers were infected with hRSV using a moi of 0.01, and at 1, 3 and 4 dpi the virus was harvested and the virus infectivity at each time of infection was determined by microplaque assay using HEp2 cell monolayers. The individual microplaques was detected by imaging of anti-P immune-stained cell monolayers using IF microscopy (Fig. 5 A and B). At 1 dpi we failed to detect microplaques, suggesting that the level of progeny virus present in the tissue culture medium at this time of infection was below the level of detection in the microplaque assay. At 3 and 4 dpi we estimated virus titers of 2.3x10^3^ pfu/ml and 3.5x10^3^ pfu/ml respectively for the virus covered from the PaKi cells (Fig. 5C). The increase in virus infectivity by 4 dpi indicated a productive RSV infection in the PaKi cell monolayers and was consistent with virus propagation in the PaKi cell line. These data also indicated that the progeny virus recovered from the PaKi cell monolayers was also able to re-infect HEp2 cells.

**Fig. 5.**
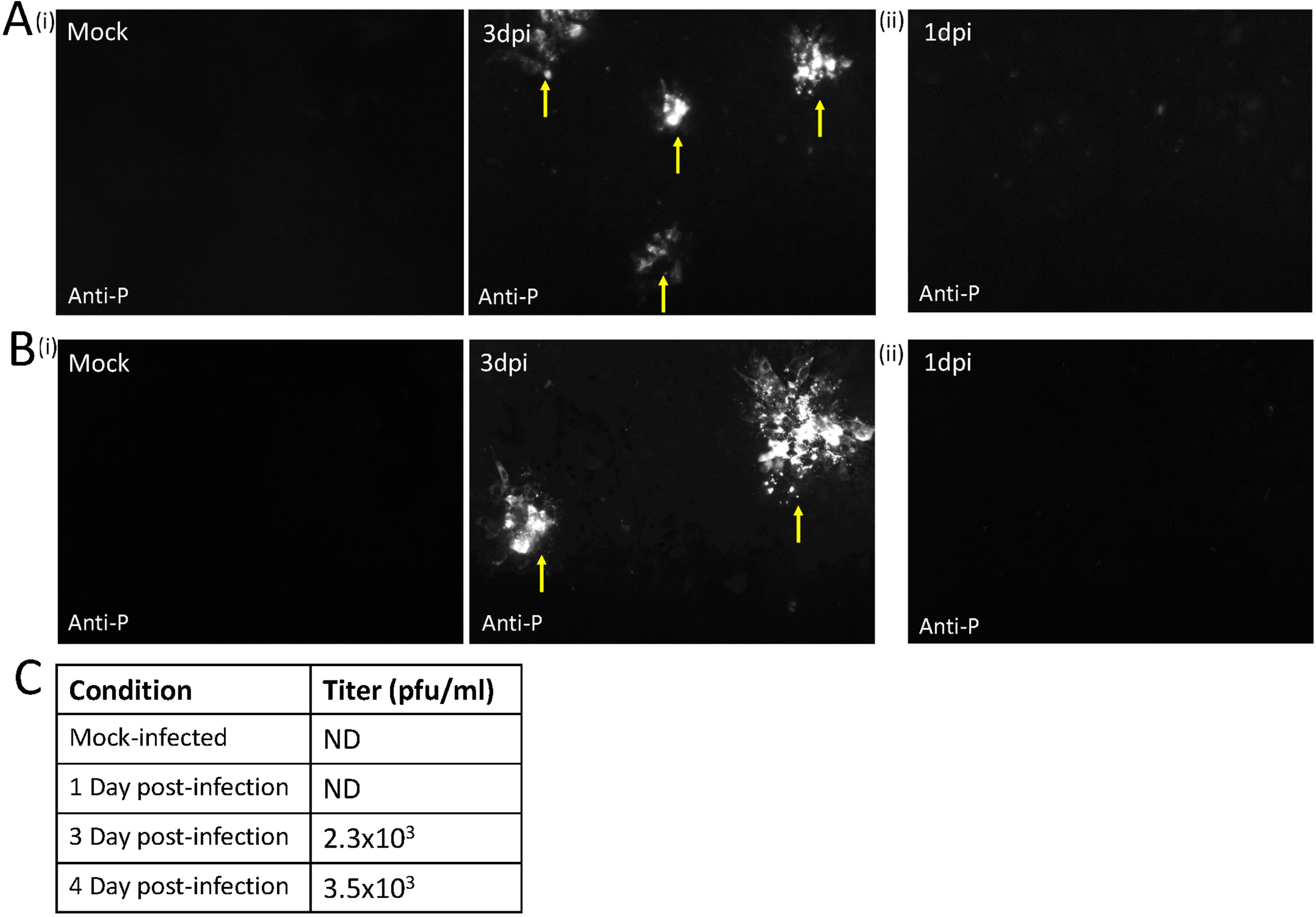
Evidence for productive RSV infection in PaKi cell monolayers. PaKi cell monolayers were mock-infected or infected with RSV using a multiplicity of infection of 0.01. **(A and B)** (i) Representative images of the plaque assay in HEp2 cells of the material harvested from the PaKi cells at 3 dpi from mock-infected and RSV-infected cells are shown. (ii) Also shown is the plaque assay of material harvested from the PaKi cells at 1 dpi. The plaques were detected using immunofluorescence microscopy to examine anti-P stained HEp2 cell monolayers: *(A)* magnification x20 objective and **(B)** magnification x40 objective, and the microplaques (yellow arrows) are highlighted. **(C)** At 1, 3 and 4 days-post infection (dpi) the virus infectivity assessed by microplaque assay is given. Data obtained from mock-infected cells at 4 dpi is also shown. ND indicates that virus infection was not detected in the HEp2 cell monolayers.

## Conclusion

Since PaKi cells were derived from bat-specific tissue, the transcriptome and proteome of the PaKi cell line is expected to reflect a bat-specific cellular environment. Our data indicates that hRSV can be efficiently propagated in PaKi cells without any prior adaptation of the virus to these cells, which suggests that the virus was able to utilize essential pro-viral host cell factors in these cells. This further suggests that PaKi cells may be a suitable cell model system with which to examine pathogen host interactions that occur in fruit bats. We can speculate that these interactions would be expected to occur at different stages of the virus replication cycle, and future work will be needed to establish the hRSV interactome in PaKi cells. It is currently unknown if hRSV can be propagated in a bat animal model, but the productive infection of hRSV in the PaKi cell line provides evidence that hRSV may also be able to replicate in tissue in bats. However, future studies will be required to determine if hRSV strains can be propagated in a fruit bat animal model.

## Materials and Methods

### Virus and cell preparation

The hRSV A2 was propagated in HEp-2 cells as described previously [18]. The PaKi cells were obtained from Prof Ling Fa Wang (DUKE-NUS, Singapore) and has been described previously [11]. The PaKi cells and HEp2 cells were maintained in DMEM (with glutamax) supplemented with 10% FCS and pen/strep at 37 ºC with 5% CO_2._

### Virus infection

Infections were performed on cells that were grown in individual wells in 24 cluster well tissue culture plate format. The low moi infections were performed in 2 well chamber slides (ThermoFischer). Infections were performed at the desired multiplicity in DMEM (with glutamax) supplemented with 2% FCS and pen/strep at 33 ºC with 5% CO_2._

### Antibodies and specific reagents

The anti-RSV (RCL3) (Novacastra Laboratories), anti-mouse and anti-rabbit IgG conjugated to Alexa 488 and Alexa555 (Molecular Probes) were purchased. The RSV monoclonal anti-G (Mab30) was obtained from Geraldine Taylor (Pirbright Institute), the polyclonal anti-F antibody was obtained from Jose Melero (Madrid, Spain), while the anti-N, anti-P and anti-M2-1 have been described previously [19].

### Microplaque titration of RSV infectivity

The recovered viral infectivity was assessed on HEp-2 cells using a microplaque assay described previously [20], but with minor modifications. Briefly, at the specific time of infection the PaKi cells were washed with PBS and the cells scraped into fresh tissue culture medium. The cell suspension was subjected to one round of freeze thaw and the cell debris removed by centrifugation (1,000 rpm/10 mins). The supernatant was diluted and used to infect near confluent HEp2 cell monolayers grown on glass coverslips. At 30 hpi the HEp-2 cell monolayers were stained using anti-P and anti-mouse IgG conjugated to Alexa 555 and the antibody-stained microplaques were counted using a Nikon eclipse 80i fluorescence microscope (Nikon ECLIPSE TE2000-U).

### Immunofluorescence microscopy

This was performed as described previously [19]. Briefly, the cells were fixed using 4% (w/v) paraformaldehyde in PBS at 4°C for 20 mins and washed using PBS at 4°C. The cells were permeabilised using 0.1% (v/v) triton X100 in PBS at 4°C for 15 mins. The cells were stained using the primary antibody and respective secondary antibody (conjugated to Alexa 488 or Alexa 555 as appropriate). The stained cells were visualized using a Nikon eclipse 80i fluorescence microscope (Nikon ECLIPSE TE2000-U) using appropriate machine settings.

### Immunoblotting analysis

The cells were washed with PBS (at 4°C), and then extracted directly into boiling mix (1% (w/v) SDS, 5% (v/v) mercaptoethanol in 20mM Tris/HCL, pH 7.5) and heated at 95°C for 2 min. The proteins were separated by 12% SDS-PAGE and transferred by Western blotting onto PVDF membrane membranes (Biorad), which were blocked with 5% skim milk in PBST (0.05% v/v Tween-20 in PBS). The PVDF membranes were then probed with the relevant primary antibody in PBST (with 5% milk) and corresponding secondary antibody conjugated to HRP. The PVDF membranes were incubated in the ECL detection system (GE Healthcare) and protein bands visualized using a the Fujifilm LAS-3000 Image system.

## Acknowledgements

We thank Dr Daniella Anderson and Professor LinFa Wang (DUKE-NUS, Singapore) for providing the PaKi cells and Aleksandra Tomczewska for technical assistance in processing of cell specimens.

